# A recombinant murine rotavirus with Nano-Luciferase expression reveals tissue tropism, replication dynamics, and virus transmission

**DOI:** 10.1101/2022.04.12.488073

**Authors:** Yinxing Zhu, Liliana Sánchez-Tacuba, Gaopeng Hou, Takahiro Kawagishi, Ningguo Feng, Harry B. Greenberg, Siyuan Ding

**Affiliations:** Department of Molecular Microbiology, Washington University School of Medicine, St. Louis, Missouri, USA; VA Palo Alto Health Care System, Department of Veterans Affairs, Palo Alto, California, USA; Department of Medicine, Division of Gastroenterology and Hepatology, Stanford School of Medicine, Stanford, California, USA; Department of Microbiology and Immunology, Stanford School of Medicine, Stanford, California, USA

**Keywords:** Rotavirus, *In vivo* imaging system, Transmission, Nano-luciferase, Tissue tropism

## Abstract

Rotaviruses (RVs) are one of the main causes of severe gastroenteritis, diarrhea, and death in children and young animals. Although suckling mice prove to be highly useful small animal models of RV infection and pathogenesis, direct visualization tools are lacking to track the temporal dynamics of RV replication and transmissibility *in vivo*. Here, we report the generation of the first recombinant murine RV that encodes a Nano-Luciferase reporter (NLuc) using a newly optimized RV reverse genetics system. The NLuc-expressing RV was replication-competent in cell culture and both infectious and virulent in neonatal mice *in vivo*. Strong luciferase signals were detected in the proximal and distal small intestines, colon, and mesenteric lymph nodes. We showed, via a noninvasive *in vivo* imaging system, that RV intestinal replication peaked at day 2 and day 5 post infection. Moreover, we successfully tracked RV transmission to uninoculated littermates as early as 3 days post infection, 1 day prior to clinically apparent diarrhea and 3 days prior to detectable fecal RV shedding in the uninoculated littermates. We also observed significantly increased viral replication in *Stat1* knockout mice that lack the host interferon signaling. Our results suggest that the NLuc RV represents a non-lethal powerful tool for the studies of tissue tropism and host and viral factors that regulate RV replication and spread, as providing a new mechanism to facilitate the testing of prophylactic and therapeutic interventions in the future.

## INTRODUCTION

Rotavirus (RV) is one of the leading causes of severe diarrhea in infants and young children. Although there are multiple safe and effective RV vaccines currently available, RV infection still results in the death of more than 128, 500 children per year [1]. Suckling mice provide a pathologically relevant small animal model for studying infection, protection, and immune responses because homologous murine RVs are a natural mouse pathogen and cause a similar diarrheal diseases as seen in human infants and many other mammalian species [2; 3]. Using this model, we and others have previously reported an important role of the type I and type III interferon (IFN) responses as well as local and systemic antibody responses in controlling RV replication in the host intestine [4; 5; 6; 7].

RV predominantly infects the host gastrointestinal tract, in particular the small intestine. However, whether RV replicates in extra-intestinal tissues such as the central nervous system, liver, and respiratory tract remains controversial [8; 9; 10; 11; 12; 13; 14]. In addition, although fecal-oral transmission is clearly the primary means of RV spread, it is technically challenging and labor intensive to follow the events of virus transmitted to naïve animals prior to the appearance of diarrheal diseases. Bioluminescent reporter systems provide extreme convenience and sensitivity to visualize intra- and inter-host viral dynamics in real time. Although fluorescent proteins and luciferase enzymes have been widely used in the studies of a variety of viral infections, including influenza virus, herpes simplex virus type 1, dengue virus, Sindbis virus, Sendai virus, and adenovirus [15; 16; 17; 18; 19; 20; 21; 22; 23], most recombinant viruses are significantly attenuated, genetically unstable, and only a few are fully applicable for *in vivo* imaging.

A plasmid-based RV reverse genetics system has recently been established and optimized by our labs and others, thereby enabling the recovery of low-titer recombinant reporter viruses and hard-to-rescue RV strains [24; 25; 26; 27]. Intragenic sequence duplications in NSP1, NSP3 and NSP5/6 gene segments have been observed in natural RV variants, leading to the production of viral proteins of unusual length and making them ideal targets to accommodate foreign gene expression [28; 29; 30]. NSP5 and NSP6 are encoded from the same gene segment, thereby introducing complications for genetic manipulation. NSP3 is expressed at higher levels than NSP1 in infected cells, rendering NSP3-based fluorescent proteins brighter and easier to detect [26]. Nano-luciferase (NLuc) is a novel bioluminescent protein and offers several advantages over the existing platforms (Firefly, Gaussia, Renilla, etc.), including enhanced stability, smaller size, and increased luminescence [31]. Hence, we take advantage of a highly efficient RV reverse genetics system that we recently developed [27] to generate a recombinant murine RV D6/2-2g strain that encodes NLuc in the RV NSP3 gene segment (rD6/2-2g-NLuc). The NLuc RV is genetically stable, replication-competent, pathogenic, and transmissible *in vivo*. Using this powerful virological tool and a well-established neonatal model of RV infection, we have begun to investigate several fundamental and important questions of RV biology including tissue tropism, replication dynamics, and virus transmission.

## RESULTS

### Generation of a recombinant NLuc-expressing murine RV

To generate rD6/2-2g-NLuc, we first constructed a T7 plasmid that expresses the NLuc reporter in the RV gene segment 7 that encodes NSP3 (pT7-NSP3-NLuc). The monomeric NLuc gene was placed downstream of the NSP3 open reading frame that is followed by a P2A self-cleaving peptide to permit separate gene expression (**Fig. 1A**). BHK-T7 cells transfected with T7-NSP3-NLuc produced the NLuc protein (**Fig. 1B**). We further confirmed by an NLuc substrate assay that strong luciferase activity was detected in T7-NSP3-NLuc-transfected cells (**Fig. 1C**). We successfully rescued the parental murine RV rD6/2-2g strain and rD6/2-2g-NLuc viruses using our optimized RV reverse genetic system [27]. NLuc expression was verified in rD6/2-2g-NLuc-infected MA104 cells (**Fig. 1D**). The identity of rD6/2-2g-NLuc was further validated by a unique electrophreotype by RNA polyacrylamide gel electrophoresis analysis (**Fig. 1E**). The edited dsRNA of RV gene segment 7 migrated slower than the wild-type gene segment 7 due to the NLuc insertion **(Fig. 1E)**. In addition, we quantified the luciferase activity in rD6/2-2g-NLuc-infected cells and found that we were able to detect signals even at the 10^5^ dilution factor **(Fig. 1F)**. Taken together, we successfully generated a murine RV NLuc reporter virus that produces robust luciferase activity in infected cells.

**Fig. 1.**
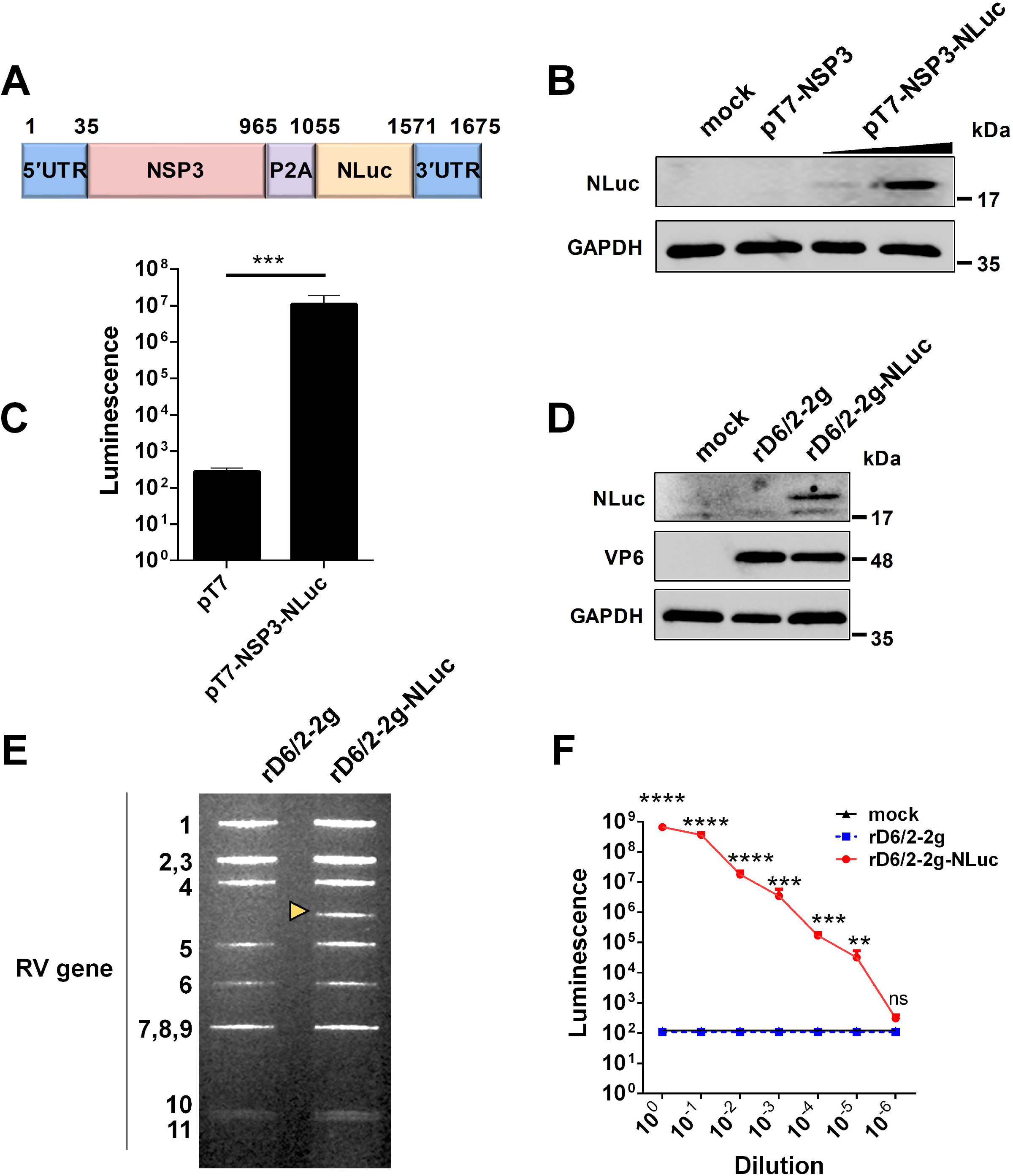
Generation and validation of a bioluminescent rD6/2-2g-NLuc. (A) A schematic diagram of a genetically engineered pT7 plasmid that encodes NLuc with nucleotide positions indicated. UTR, untranslated region; P2A, self-cleaving P2A peptide gene of porcine teschovires-1. (B) BHK-T7 cells were transfected with pT7-NSP3 and increasing amounts of pT7-NSP3-NLuc for 48 hours, and cell lysates were analyzed by western blot. (C) BHK-T7 cells were transfected with pT7 or pT7-NSP3-NLuc plasmids for 48 hours. The luciferase activity was determined by Nano-Glo® luciferase assay. Data are presented as the average of three experiments and error bars indicate standard error of the mean (SEM) (Student t test; *** P < 0.001). (D) MA104 cells were infected with rD6/2-2g and rD6/2-2g-NLuc viruses (MOI=0.1) for 24 hours, and cell lysates were analyzed by western blot. (E) dsRNA profiles. Viral RNA was extracted from sucrose cushion-concentrated virus, separated on a 10% polyacrylamide gel, and then stained with ethidium bromide. The dsRNA segment numbers are indicated and the position of the engineered segment 7 is marked with a yellow arrowhead. (F) Luciferase activity of rD6/2-2g and rD6/2-2g-NLuc. MA104 cells were infected with 10-fold serially diluted rD6/2-2g or rD6/2-2g-NLuc. Cells were harvested at 48 hpi and the luciferase activity was determined by Nano-Glo® luciferase assay. Results are expressed as the mean luminesence of triplicates and error bars show the SEM (one-way ANOVA with Dunnett’s test; ns, not significant, * P < 0.05, ** P < 0.01, *** P < 0.001, **** P < 0.0001).

### Characterization of rD6/2-2g-NLuc replication *in vitro*

We next sought to determine the replication kinetics of rD6/2-2g-NLuc as compared to the parental rD6/2-2g *in vitro*. Despite slightly lower intracellular mRNA levels and virus titers than those of rD6/2-2g at 24, 48, and 72 hours post infection (hpi) **(Fig. 2A and 2B)**, rD6/2-2g-NLuc replicated well in MA104 cells and produced substantial cytopathic effects (data not shown). The plaque size of rD6/2-2g-NLuc was approximately half of that of rD6/2-2g **(Fig. 2C)**. We performed serial passage of rD6/2-2g-NLuc in MA104 cells to assess the genetic stability. Importantly, luminescence was still highly detectable after 8 passages and we observed no loss of luciferase signals over time **(Fig. 2D)**, suggesting that the NLuc gene was functionally maintained in the viral genome and that the reporter virus is infectious and stable *in vitro*.

**Fig. 2.**
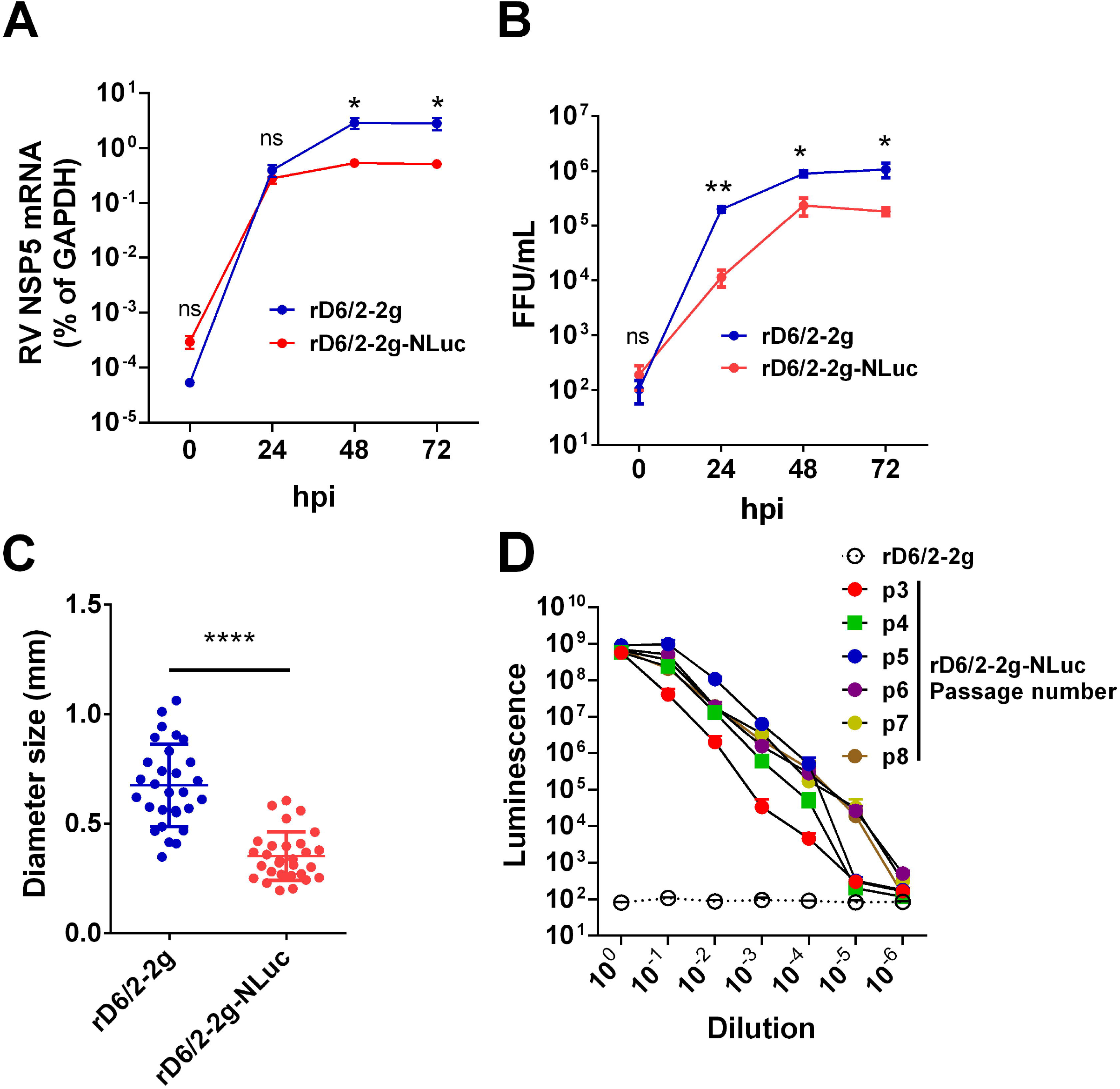
Growth kinetics of bioluminescent rD6/2-2g-NLuc in MA104 cells. (A) MA104 cells were infected with rD6/2-2g or rD6/2-2g-NLuc (MOI=0.01) in the presence of trypsin (0.5Lμg/ml) and harvested at the indicated time points. The viral mRNA level was determined by RT-qPCR assay and normalized to that of GAPDH. Data are the average of three experiments, error bars indicate SEM (two-way ANOVA test; ns, not significant, * P < 0.05, ** P < 0.01). (B) Multi-step growth curves of rD6/2-2g-NLuc. MA104 cells were infected with rD6/2-2g or rD6/2-2g-NLuc (MOI=0.01) in the presence of trypsin (0.5Lμg/ml) and harvested at the indicated time points. The viral titers were determined by an immunoperoxidase focus-forming assay. Data are the average of three experiments, error bars indicate SEM (two-way ANOVA test; ns, not significant, * P < 0.05, ** P < 0.01). (C) Plaque formation of rD6/2-2g-NLuc. Plaques were generated on MA104 monolayers and detected by crystal violet staining at 7 dpi. The diameter of at least 25 randomly selected plaques from 2 independent plaque assays was measured by a bright-field microscope. Error bars indicate SEM (Student t test; **** P < 0.0001). (D) Functional stability of luciferase activity in rD6/2-2g-NLuc after sequential passage. rD6/2-2g-NLuc was sequentially passaged in MA104 cells. The luciferase activity for passages 3-8 was determined by Nano-Glo® luciferase assay as described. Results are expressed as the mean luminesce of duplicates. Error bars show SEM. Luminescence from NLuc substrate from MA104 cells infected with rD6/2-2g were plotted as a reference.

### Tissue tropism of rD6/2-2g-NLuc *in vivo*

To leverage the high sensitivity of NLuc and investigate RV tissue tropism, we orally inoculated five-day-old 129sv pups with 1.3×10^6^ foci forming units (FFUs) of rD6/2-2g-NLuc. We observed 100% diarrheal development in infected pups at 1 day post infection (dpi) **(Fig. 3A)**. The diarrhea occurrence remained more than 50% from 2 to 5 dpi **(Fig. 3A)**. We euthanized one mouse on each day and harvested different organs to measure luciferase activities. As expected, we found strong luciferase signals throughout the lower gastrointestinal tract. We detected more robust activity in the distal small intestine (SI) than proximal SI (**Fig. 3B and 3C**). We also detected high NLuc activity in the colon and the mesenteric lymph nodes (**Fig. 3D and 3E**), suggestive of active RV replication at these sites. On the other hand, the pancreas and the liver had weak to non-detectable signals **(Fig. 3F and 3G)**. These results suggest that murine RV primarily targets the lower gastrointestinal tract (SI and colon) and does not actively replicate in extra-intestinal organs such as the liver.

**Fig. 3.**
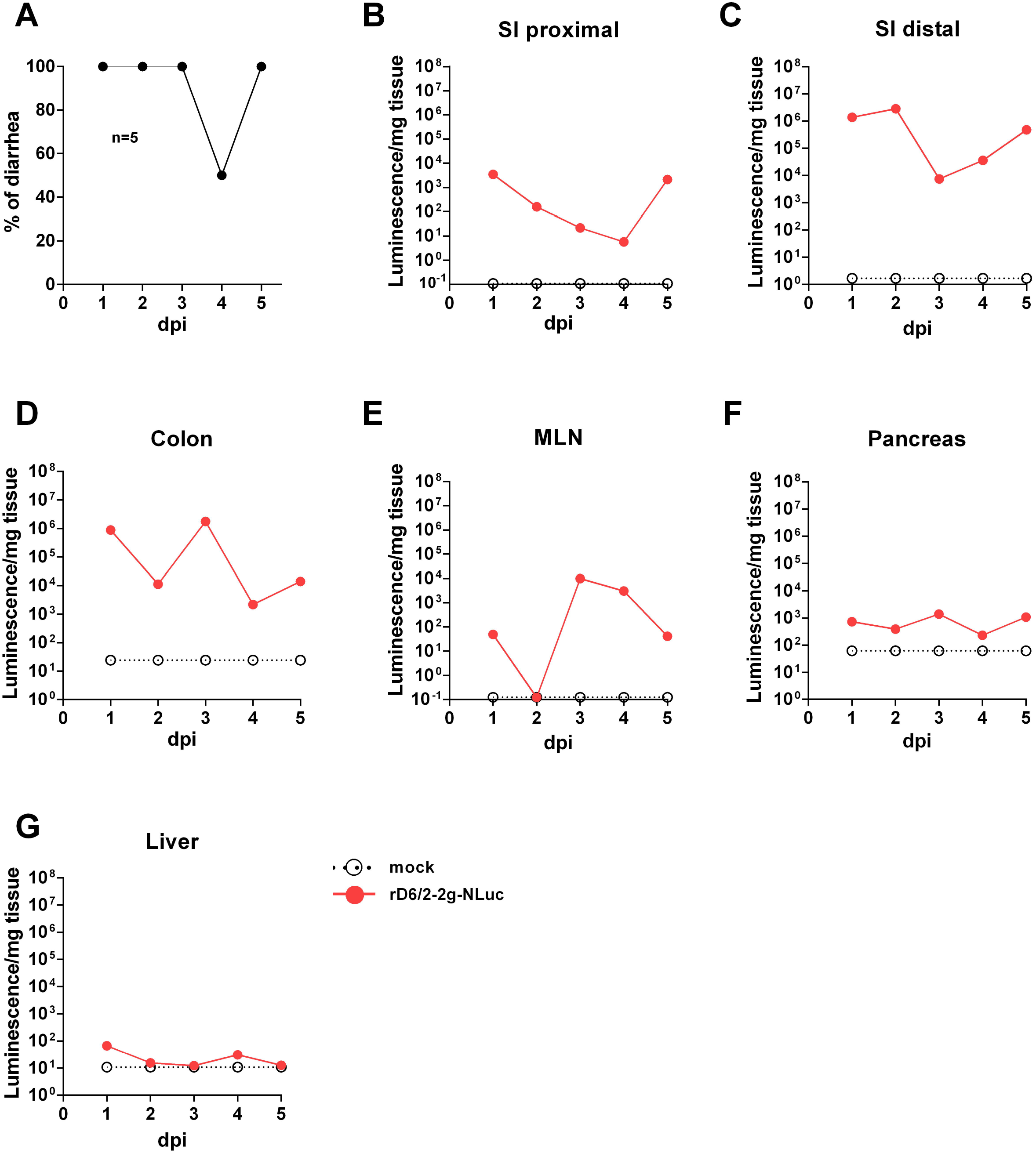
Bioluminescence of rD6/2-2g-NLuc in the intestines and the systemic sites in wild-type 129sv mice. (A) Five-day-old wild-type 129sv pups (n=5) were orally infected with 1.3□×□10^6^ FFUs of rD6/2-2g-NLuc and diarrhea was monitored till 5 days post infection. (B-G) Five-day-old wild-type 129sv pups were orally infected with 1.3□×□10^6^ FFUs of rD6/2-2g-NLuc, then euthanized at indicated days post infection. Bioluminescence from indicated tissue homogenates was determined by Nano-Glo® luciferase assay. Luminescence from NLuc substrate of uninfected mice tissues were plotted as a reference.

### Infectivity and pathogenicity of rD6/2-2g-NLuc *in vivo*

To investigate whether we can use rD6/2-2g-NLuc for studies of intestinal RV infection, we inoculated five-day-old 129sv mice with a low inoculum (3.5□×□10^3^ FFUs) of rD6/2-2g-NLuc via oral gavage. We observed that 50% of mice developed diarrhea at 1 dpi and about 80% developed diarrhea by 2 dpi **(Fig. 4A)**. We found high levels of fecal shedding of infectious RVs from 4 to 10 dpi **(Fig. 4B)**. Importantly, we recorded the bioluminescence signals from day 0 to day 12 post infection and observed strong luciferase in the abdominal cavity as early as 1 dpi using the *in vivo* imaging system (IVIS) **(Fig. 4C)**. The luminescence intensity was over to 10^6^ p/sec/cm^2^/sr and remained high until 7 dpi **(Fig. 4D)**. These results demonstrate that our reporter virus provides extreme sensitivity and temporal resolution of intra-intestinal RV infection multiple days prior to the detection of RV shedding in the fecal specimens.

**Fig. 4.**
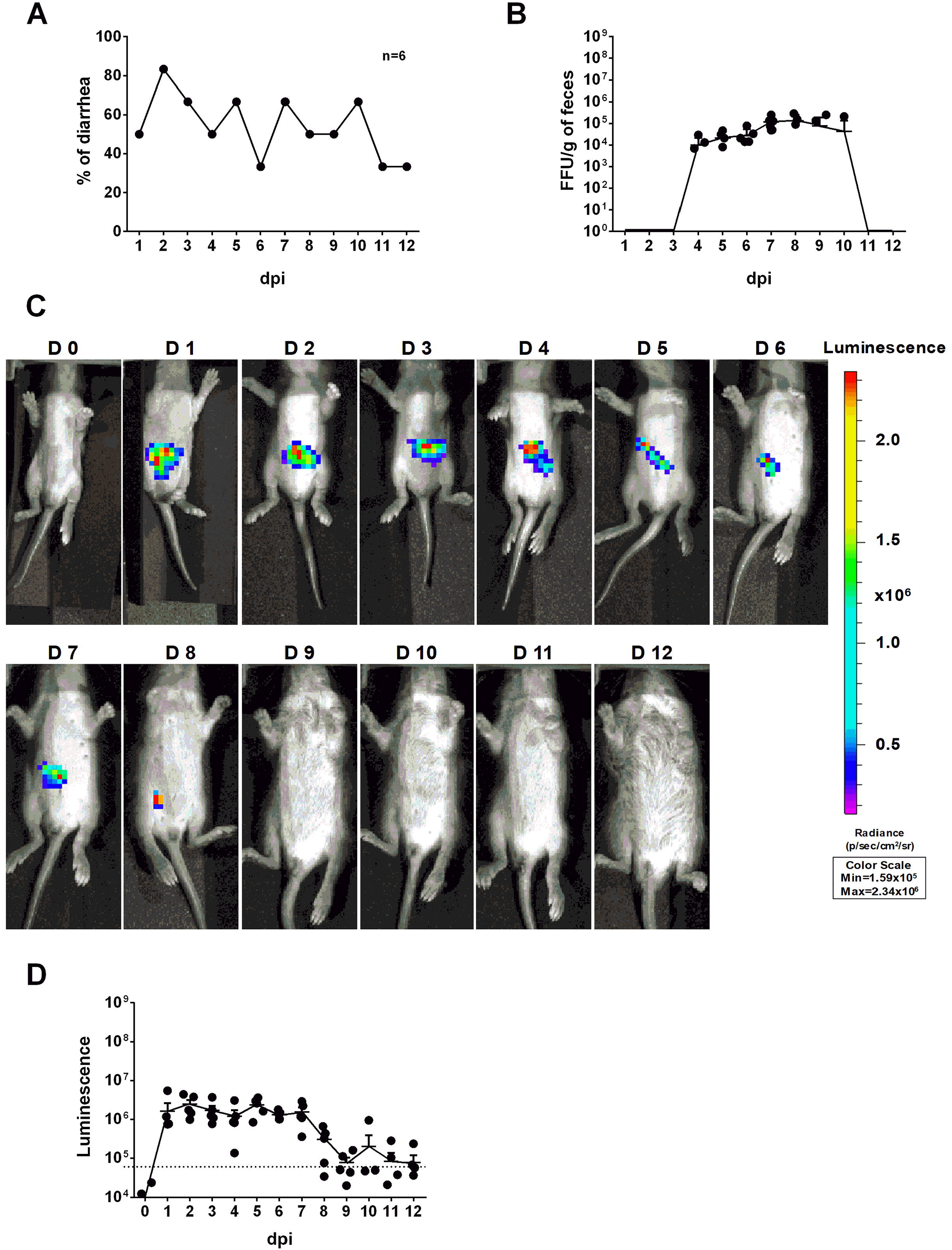
Infectivity and pathogenicity of rD6/2-2g-NLuc *in vivo*. (A) Five-day-old 129sv mice (n=6) were orally inoculated with 3.5□×□10^3^ FFUs of rD6/2-2g-NLuc. The diarrhea rate was monitored from 1 to 12 days post infection. (B) Viral shedding in stool samples was detected by an FFU assay and normalized to the feces weight. (C) Representative images of rD6/2-2g-NLuc infected pups (1 to 12 days). The bioluminescent signal is expressed in photons per second per square centimeter per steradian (p/sec/cm^2^/sr). (D) Quantification of the luminescence in (C). The dashed line indicates the upper limit of detection.

### Characterization of RV transmission by IVIS

To further quantitatively track RV transmission, an important but under-studied aspect of RV biology, we co-housed 6 infected and 6 uninfected littermates in the same cage. Compared to the RV-inoculated mice (**Fig. 4A**), diarrhea was first observed in the naïve animals at 4 dpi and reached over 80% at 7 dpi **(Fig. 5A)**. We also quantified RV fecal shedding by an FFU assay. The originally uninoculated mice had detectable virus shedding briefly between 6 to 8 dpi **(Fig. 5B)**, albeit at a similar level as the infected mice (**Fig. 4B**). Remarkably, we observed strong luminescence as early as 3 dpi **(Fig. 5C and 5D)**, preceding the first appearance of clinical symptoms at 4 dpi and fecal shedding at 6 dpi. These data indicate that RV transmission readily occurred 3 days after co-housing and that rD6/2-2g-NLuc is a highly sensitive and convenient tool for following RV infection and spread in real time *in vivo*.

**Fig. 5.**
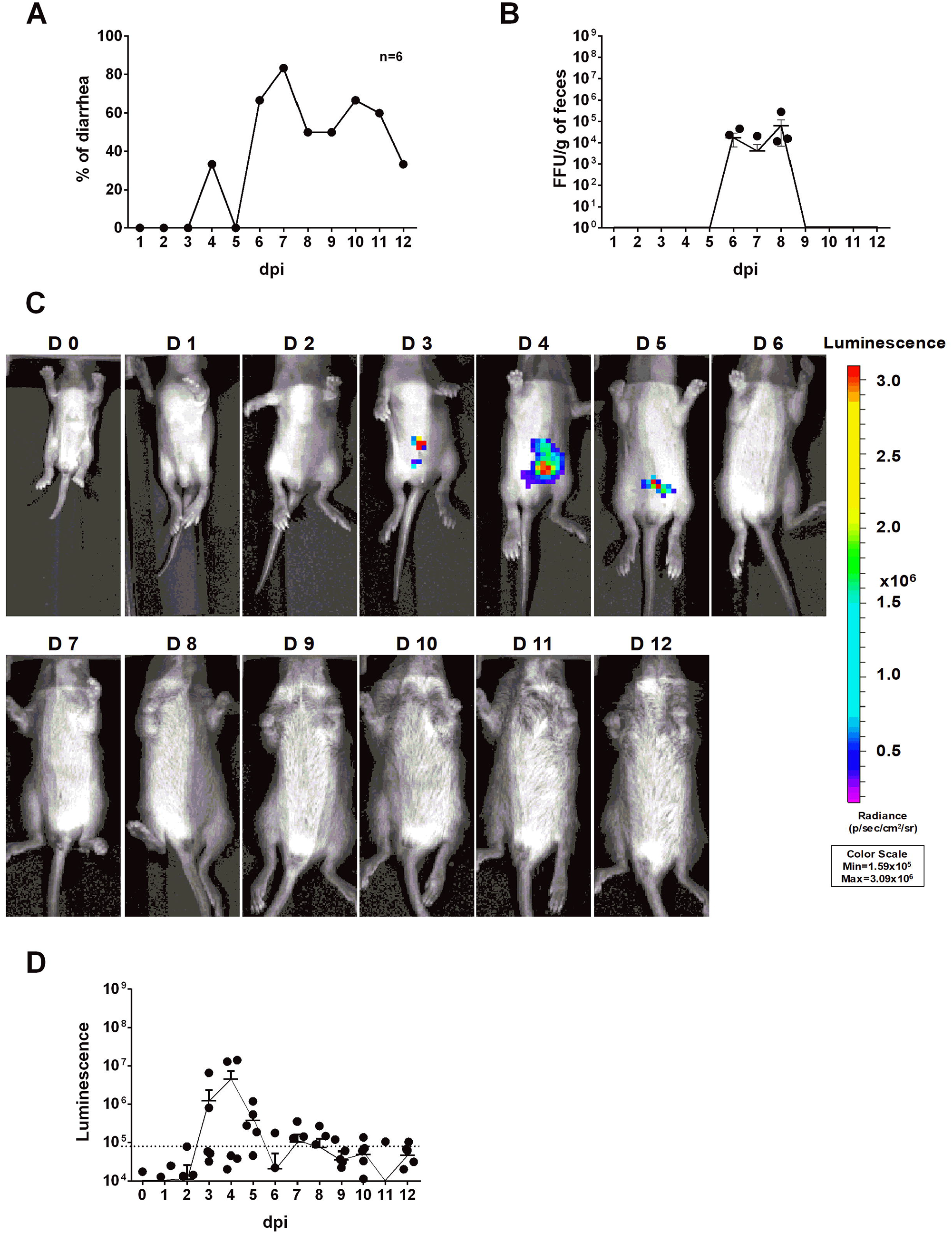
Transmission rD6/2-2g-NLuc *in vivo*. (A) Five-day-old 129sv mice were co-housed with 6 infected (3.5□×□10^3^ FFUs of rD6/2-2g-NLuc) and 6 uninfected littermates in the same cage. The diarrhea rate was monitored from 1 to 12 days post infection. (B) Viral shedding in stool samples was detected by an FFU assay and normalized to the feces weight. (C) Representative images of naive pups (1 to 12 days). The bioluminescent signal is expressed in photons per second per square centimeter per steradian (p/sec/cm^2^/sr). (D) Quantification of the luminescence in (C). The dashed line indicates the upper limit of detection.

### RV infection of *Stat1* knockout mice

To determine whether IVIS represents an ideal system to study the role of host factors in RV intestinal replication, which is enhanced in immunodeficient mice, we orally infected five-day-old *Stat1* knockout (KO) mice with 3.5□×□10^3^ FFUs of rD6/2-2g-NLuc, at the same dose as in wild-type 129sv mice **(Fig. 4)**. We observed that about 30% of mice developed diarrhea at 1 dpi and 100% developed diarrhea from 2 until 6 dpi **(Fig. 6A)**. As expected, *Stat1* KO pups had high levels of fecal shedding of infectious virus particles at 1 to 3 dpi **(Fig. 6B)**, much earlier than that observed in the wild-type animals **(Fig. 4B)**. Moreover, IVIS revealed that the luminescence intensity was significantly increased (approximately 10-fold higher, up to 10^7^ p/sec/cm^2^/sr) with the lack of host interferon signaling **(Fig. 6C-E)**. Collectively, these results demonstrate the utility and effectiveness of rD6/2-2g-NLuc in objectively reflecting RV replication and studying host immunity *in vivo*.

**Fig. 6.**
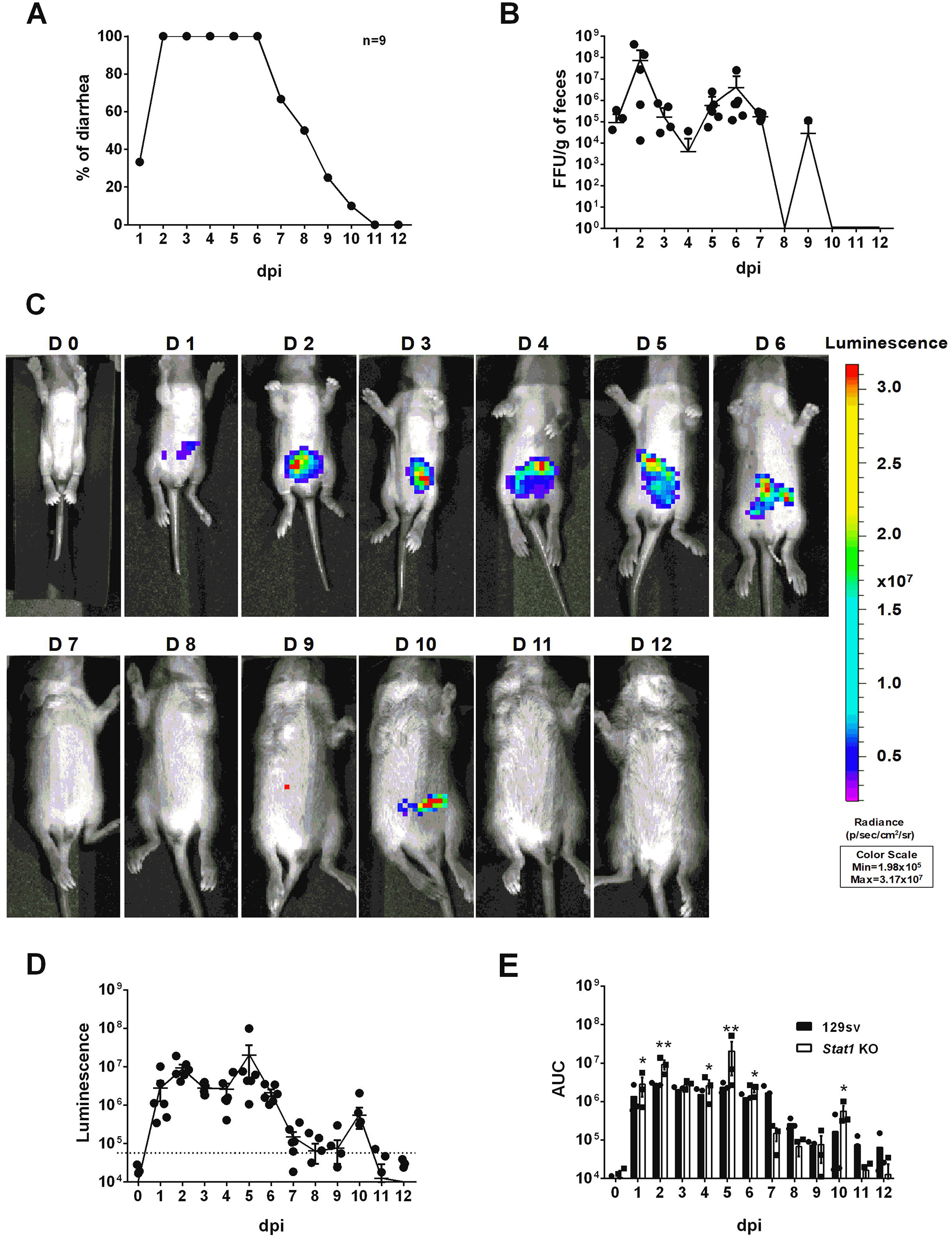
Characterization of rD6/2-2g-NLuc infection in *Stat1* KO 129sv mice. (A) Five-day-old *Stat1* KO 129sv mice (n=9) were orally inoculated with 3.5×□10^3^ FFUs of rD6/2-2g-NLuc. The diarrhea rate was monitored from 1 to 12 days post infection. (B) Viral shedding in stool samples was detected by an FFU assay and normalized by to feces weight. (C) Representative images of rD6/2-2g-NLuc infected *Stat1* KO pups (1 to 12 days). The bioluminescent signal is expressed in photons per second per square centimeter per steradian (p/sec/cm^2^/sr). (D) Quantification of the luminescence in (C). The dashed line indicates the upper limit of detection. (E) Statistical analysis of area under the curve (AUC) comparing data in Fig. 4D and 6D. Error bars show the SEM (one-way ANOVA test; ns, not significant, * P < 0.05, ** P < 0.01).

## DISCUSSION

Reporter viruses prove to be important tools for visualizing and monitoring viral replication dynamics *in vitro* and *in vivo*. Although the plasmid-based RV reverse genetics system was reported in 2017 and several fluorescent and luminescent protein-encoding RVs (primarily in the backbone of simian RV SA11 strain) have been reported [24; 25; 26], murine viruses are difficult to rescue, precluding further manipulation and heterologous expression of foreign genes. In this study, we take advantage of a more efficient RV reverse genetics system that we recently developed [27] and generate a murine rD6/2-2g-NLuc strain. This virus was functionally stable even after 8 passages **(Fig. 2D)**. The combinatorial use of rD6/2-2g-NLuc reporter virus and IVIS enabled the detection of RV replication in different organ systems (**Fig. 3**). It is noteworthy that we found strong luciferase signals in the colon (**Fig. 3D**). There is controversy in the literature regarding RV infection of the large intestine including cecum and colon [32; 33; 34; 35; 36; 37]. With the limitation that we cannot distinguish real infection of colon epithelium from NLuc activities from infected shed but still alive cells, our data suggest a strong possibility that RV also replicates in the colon. It is also interesting that we transiently detected strong luciferase signals in the mesenteric lymph node (**Fig. 3E**), composed predominantly of hematopoietic cells. Furthermore, our system allowed us to assess RV transmission to uninoculated co-caged littermates **(Fig. 5)** and the effect of host factors and signaling pathways on RV intestinal replication *in vivo* **(Fig. 6)**, which is easily extendable to the role of other host innate and adaptive antiviral signaling in RV infection, pathogenesis, and transmission.

Given the modular nature and small size of the NLuc reporter construct, this approach is broadly applicable to the studies of other RV isolates and other enteric viruses (murine norovirus, enterovirus D68, etc.). The replication of simian RVs is severely limited in immunocompetent suckling mice. To that end, we can rescue NLuc reporter in the backbone of simian RV RRV strain, which we expect to be attenuated in 129sv mice but to cause a lethal biliary disease in *Stat1* deficient or intra-peritoneally inoculated newborn mice [38]. We can apply traditional virological approaches (reassortment with gene swapping and/or deletion) to examine the relative contribution of individual RV gene product in intestinal replication and transmission. Another interesting aspect is that the host genetic background dictates RV pathogenicity. Compared to 129sv pups, diarrhea in C57Bl/6 mice is highly attenuated. Thus, one could use NLuc virus to desegregate RV replication from diseases and help dissect the role of RV-encoded products in this process.

Finally, reporter viruses have emerged as powerful tools in small-molecule compound screening [39; 40], antibody identification [16], and vaccine efficacy analysis [41]. We envision that our NLuc reporter RV and IVIS will provide a rapid, non-lethal and real-time quantitative means to assess viral replication, spread and facilitate the rationale design and development of novel antiviral therapeutics and new-generation safe and efficacious RV vaccines, to be tested in pre-clinical small animal models.

## MATERIAL AND METHOD

### Cell culture and viruses

MA104 cells (ATCC CRL-2378) were cultured in Medium 199 (M199, Sigma-Aldrich) supplemented with 10% heat-inactivated fetal bovine serum (FBS), 100 I.U. penicillin/ml, 100 µg/ml streptomycin and 0.292 mg/ml L-glutamine (complete medium). The BHK-T7 cell line [42] was provided by Dr. Ursula Buchholz (Laboratory of Infectious Diseases, NIAID, NIH, USA) and cultured in completed DMEM supplemented with 0.2 μg/ml of G-418 (Promega). MA104 N*V cells were cultured in complete M199 in the presence of 3 μg/ml puromycin and 3 μg/ml of blasticidin (InvivoGen, San Diego, CA).

The rRV strains used in this study include rD6/2-2g and rD6/2-2g-NLuc and were propagated in MA104 cells. Prior to infection, all RV inocula were activated with 5 μg/ml of trypsin (Gibco Life Technologies, Carlsbad, CA) for 30 min at 37 °C.

### Plasmid construction

The murine D6/2 rescue plasmids: pT7-D6/2-VP2, pT7-D6/2-VP3, pT7-D6/2-VP4, pT7-D6/2-VP6, pT7-D6/2-VP7, pT7-D6/2-NSP1, pT7-D6/2-NSP2, pT7-D6/2-NSP3, and pT7-D6/2-NSP5 were prepared as described previously [27] while pT7-SA11-VP1 and pT7-SA11-NSP4 were originally made by Dr. Takeshi Kobayashi (Research Institute for Microbial Diseases, Osaka University, Japan) [24] and obtained from Addgene. The C3P3-G1 plasmid [43] was kindly provided by Dr. Philippe H Jaïs. To generate pT7-D6/2-NSP3-NLuc, which encodes a full-length NLuc gene (GenBank: KM359774.1) and the self-cleaving P2A peptide gene of porcine teschovires-1, the P2A-NLuc gene cassette was amplified by PCR and inserted between nucleotides in the NSP3 gene via Gibson assembly (NEBuilder HiFi DNA Assembly kit). Purification of all the plasmids was performed using QIAGEN Plasmid Maxiprep kit per the manufacturer’s instructions.

### Generation of recombinant rotaviruses

rD6/2-2g was generated using the following pT7 plasmids: pT7-SA11-VP1 and -NSP4, pT7-D6/2-VP2, -VP3, -VP4, -VP6, -VP7, -NSP1, -NSP2, -NSP3 and -NSP5 according to the optimized entirely plasmid-based RG system [27]. The pT7-D6/2-NSP3 plasmid was replaced by the pT7-D6/2-NSP3-NLuc to generate rD6/2-2g-NLuc. The rescued rRVs were propagated for two passages in MA104 cells in a 6-well plate, and then both rRVs were plaque purified twice in MA104 cells.

### Western blot

BHK-T7 cells were transfected with 1µg pT7 vector or 1 and 2 µg pT7-NSP3-NLuc plasmids for 48 hours and MA104 cells were infected by rD6/2-2g or rD6/2-2g-NLuc at an MOI of 0.1 for 24 h. Then, cells were washed twice with ice-cold phosphate-buffered saline (PBS; Thermo Scientific) and lysed in RIPA buffer (150□mM NaCl, 1.0% IGEPAL CA-630, 0.5% sodium deoxycholate, 0.1% SDS, 50□mM Tris, pH 8.0; Sigma-Aldrich) supplemented with 1× protease inhibitor cocktail (Thermo Scientific) for 30□min at 4°C. After that, cell debris was removed by centrifugation at 12,000□×□g for 10□min at 4°C. Samples were resolved in precast SDS-PAGE gels (4 to 15%; Bio-Rad) and transferred to a nitrocellulose membrane (0.45□μm; Bio-Rad). The membrane was incubated with blocking buffer (5% bovine serum albumin [BSA] diluted in PBS supplemented with 0.1% Tween 20) for 1 h at room temperature. Then, the membrane was incubated with anti-NLuc rabbit monoclonal antibody (Promega; catalog no. N7000; 1 µg/ml) diluted in SuperBlock blocking buffer, 4 °C overnight, anti-RV VP6 mouse monoclonal antibody (Santa Cruz Biotechnology; sc-101363; 1:1,000), and anti-glyceraldehyde-3-phosphate dehydrogenase (GAPDH) rabbit monoclonal antibody (CST; catalog no. 2118; 1:1,000), followed by incubation with anti-mouse IgG (CST; catalog no. 7076; 1:5,000) or anti-rabbit IgG (CST; catalog no. 7074; 1:5,000) horseradish peroxidase (HRP)-linked antibodies. The antigen-antibody complex was detected using Clarity Western ECL substrate (Bio-Rad) and the ChemiDoc MP imaging system according to the manufacturer’s manuals.

### RT-qPCR

The total RNA of the MA104 cells infection with recombinant rD6/2-2g and rD6/2-2g-NLuc virus was extracted by TRIzol. Total RNA was reverse transcribed to cDNA using a high-capacity cDNA reverse transcription kit with RNase inhibitor (Applied Biosystems) according to the user guide. Briefly, 0.8□μg of RNA, 2□μl of 10× reverse transcription (RT) buffer, 0.8□μl of 100□mM deoxynucleoside triphosphate (dNTP) mix, 2□μl of RT random primers, 0.1□μl of RNase inhibitor, 0.1□μl of MultiScribe reverse transcriptase, and a flexible amount of nuclease-free H_2_O were added to the 20 μl reaction mixture. The reverse transcription thermocycling program was set at 25°C for 10□min, 37°C for 2 h, and 85°C for 5□min. The expression level of housekeeping gene GAPDH was quantified by 2× SYBR green master mix (Applied Biosystems), and NSP5 was measured by 2× TaqMan Fast Advanced master mix (Applied Biosystems). The primers used in this study were as follows: human GAPDH forward primer, 5′-GGAGCGAGATCCCTCCAAAAT-3′, and reverse primer, 5′-GGCTGTTGTCATACTTCTCATGG-3′; and NSP5 forward primer, 5′-CTGCTTC AAACGATCCACTCAC-3′, reverse primer, 5′-TGAATCCATAGACACGCC-3′, and probe, 5′-CY5/TCAAATGCAGTTAAGACAAATGCAGACGCT/IABRQSP-3′. The y axis stands for the percentage of NSP5 mRNA levels relative to GAPDH levels.

### Plaque assay

Activated virus samples were serially diluted 10-fold and added to monolayers of MA104 cells for 1 h at 37°C. Inocula were removed and replaced with 0.1% (w/v) agarose (SeaKem® ME Agarose. Lonza) in FBS-free M199 supplement with 0.5 μg/ml of trypsin. Cultures were incubated for 7 days at 37°C in a 5% CO2 incubator. Random plaques were picked by pushing the 200 μl tip through the overlay agarose, and then were propagated in MA104 cells as described above. To quantify the plaque diameter, cultures at 7 dpi were fixed with 10% formaldehyde and stained with 1% crystal violet (Sigma-Aldrich). The diameter of at least 25 randomly selected plaques from 2 independent plaque assays was recorded using an ECHO microscope and then, diameters were measured with the annotation tool of the microscope.

### Focus-Forming assay

Activated virus samples from cell culture or mouse stool specimens were serially diluted 2- or 10-fold and added to confluent monolayers of MA104 cells seeded in 96-well plates for 1 h at 37°C. Inocula were removed and replaced with M199 serum-free and then incubated for 16 to18 h at 37 °C. Cells were then fixed with 10% paraformaldehyde and permeabilized with 1% Tween 20. Cells were incubated with rabbit hyperimmune serum to rotavirus (anti-DLPs) produced in our laboratory and anti-rabbit HRP-linked secondary antibody. Viral foci were stained with 3-amino-9-ethylcarbazole (AEC substrate kit. Vector Laboratories) per manufacturer’s instructions and enumerated visually.

### Luciferase assay

MA104 cells seeded in 96-well plates were infected with 50 µL of 10-fold serial dilution of rRVs at 37°C for 48 h and freeze-thawed 2 times before 50 µL/well of Nano-Glo Luciferase Assay Reagent (Promega) was added per manufacturer’s instructions. After 5 minutes incubation at room temperature, relative luminosity units were measured using a 20/20n Luminometer (Turner Biosystems). 100 μL of mouse tissues homogenates were mixed with 50 μL of Nano-Glo working substrate solution and processed as described above.

### Purification of RV particles by sucrose gradient centrifugation

rRVs were concentrated by pelleting through a sucrose cushion as described [44]. Briefly, MA104 grown in 12-well plate were infected and harvested 72 h post infection (hpi), the viral lysates were freeze-thawed three times, and viral particles concentrated by ultracentrifugation for 1 h at 30,000 rpm at 4 °C. Viral pellets were resuspended in TNC buffer (10 mM Tris-HCl [pH 7.5], 140 mM NaCl, 10 mM CaCl2), extracted with genetron and the aqueous phase pelleted through a 40% sucrose cushion by centrifugation for 1 h at 30,000 rpm at 4°C. The pelleted rRV was resuspended with 1 mL of PBS with Ca^2+^ and Mg^2+^ and this suspension was used to perform mice infections or to obtain genomic dsRNA profiles.

### Electrophoresis of viral dsRNA genomes

Viral dsRNAs were extracted from sucrose cushion-concentrated rRVs with TRIzol (Invitrogen) according to the manufacturer’s protocol and then mixed with Gel Loading Dye, Purple (6x), no SDS (NEB). Samples were subjected to PAGE (10%) for 2h 30 min at 180V and then stained with ethidium bromide (0.1 µg/mL) for 10 minutes and visualized by the gel documentation system (Axygen).

### Mice infection and phenotypic characterization

Wild-type 129sv and *Stat1* KO mice were purchased from the Jackson Laboratory and Taconic Biosciences and bred locally at the Washington University in St. Louis (WUSTL) CSRB vivarium. Wild type 129sv mice were originally purchased from the Jackson Laboratory and maintained in-house in a breeding colony. 5-day-old pups were orally inoculated with rD6/2-2g-NLuc (1.3□×□10^6^ FFU) or PBS. Diarrhea was scored as previously described (Broome et al). On the indicated day animals were sacrificed and small intestine, colon, mesenteric lymph node, pancreas, and liver were collected, weighed, homogenized in PBS with Ca^2+^ and Mg^2+^ and clarified by centrifugation. Homogenized tissues were subjected to Luciferase activity. Proximal and distal small intestines samples were collected: proximal samples were collected at about 2-3 cm from the pyloric sphincter while distal were collected at about 0.5 cm from the caecum.

### IVIS

Wild-type 129sv and *Stat1* KO mice were purchased from the Jackson Laboratory and Taconic Biosciences and bred locally at the Washington University in St. Louis (WUSTL) CSRB vivarium. Five-day-old suckling pups were orally infected with rD6/2-2g-NLuc (3.5□×□10^3^ FFU). Diarrhea was evaluated from day 1 to day 12 post infection. To perform IVIS, we firstly weighted the mice, and oral gavage Nano-Glo™ substrate (1/20 dilution in PBS; to make sure 50 µL per mouse, 1/25-1/57 dilution in PBS) for 3.5 hours and then performed IVIS (exposure time: 1 second) by using the IVIS Spectrum BL.

### Statistical analysis

All statistical tests were performed as described in the indicated figure legends using Prism 9.0. Statistical significance was determined using a one-way ANOVA when comparing three or more groups. When comparing two groups, a Mann-Whitney test and student t test were performed. The number of an independent experiment performed is indicated in the relevant figure legends.

## Acknowledgement

We thank the members of the Ding lab for helpful discussion of the project. We appreciate Drs. Nathan J. Meade and Kenneth H. Mellits for kindly sharing the MA104-N*V cells and Dr. Philippe H. Jais for sharing the C3P3-G1 plasmid. We thank Drs. Suzanne M. Hickerson and Stephen M. Beverley for the IVIS training.

## REFERENCES

[1] C. Troeger, I.A. Khalil, P.C. Rao, S.J. Cao, B.F. Blacker, T. Ahmed, G. Armah, J.E. Bines, T.G. Brewer, D.V. Colombara, G. Kang, B.D. Kirkpatrick, C.D. Kirkwood, J.M. Mwenda, U.D. Parashar, W.A. Petri, M.S. Riddle, A.D. Steele, R.L. Thompson, J.L. Walson, J.W. Sanders, A.H. Mokdad, C.J.L. Murray, S.I. Hay, and R.C. Reiner, Rotavirus Vaccination and the Global Burden of Rotavirus Diarrhea Among Children Younger Than 5 Years. Jama Pediatr 172 (2018) 958–965.

[2] M.A. Franco, J. Angel, and H.B. Greenberg, Immunity and correlates of protection for rotavirus vaccines. Vaccine 24 (2006) 2718–31.

[3] J.W. Burns, A.A. Krishnaney, P.T. Vo, R.V. Rouse, L.J. Anderson, and H.B. Greenberg, Analyses of homologous rotavirus infection in the mouse model. Virology 207 (1995) 143–53.

[4] N. Feng, L.L. Yasukawa, A. Sen, and H.B. Greenberg, Permissive replication of homologous murine rotavirus in the mouse intestine is primarily regulated by VP4 and NSP1. Journal of virology 87 (2013) 8307–16.

[5] J.D. Lin, N. Feng, A. Sen, M. Balan, H.C. Tseng, C. McElrath, S.V. Smirnov, J. Peng, L.L. Yasukawa, R.K. Durbin, J.E. Durbin, H.B. Greenberg, and S.V. Kotenko, Distinct Roles of Type I and Type III Interferons in Intestinal Immunity to Homologous and Heterologous Rotavirus Infections. PLoS pathogens 12 (2016) e1005600.

[6] N.G. Feng, M.C. Jaimes, N.H. Lazarus, D. Monak, C.Q. Zhang, E.C. Butcher, and H.B. Greenberg, Redundant role of chemokines CCL25/TECK and CCL28/MEC in IgA(+) plasmablast recruitment to the intestinal lamina propria after rotavirus infection. J Immunol 176 (2006) 5749–5759.

[7] E.M. Deal, K. Lahl, C.F. Narvaez, E.C. Butcher, and H.B. Greenberg, Plasmacytoid dendritic cells promote rotavirus-induced human and murine B cell responses. J Clin Invest 123 (2013) 2464–2474.

[8] M. Santosham, R.H. Yolken, E. Quiroz, L. Dillman, G. Oro, W.C. Reeves, and R.B. Sack, Detection of rotavirus in respiratory secretions of children with pneumonia. The Journal of pediatrics 103 (1983) 583–5.

[9] M. Fragoso, A. Kumar, and D.L. Murray, Rotavirus in nasopharyngeal secretions of children with upper respiratory tract infections. Diagnostic microbiology and infectious disease 4 (1986) 87–8.

[10] B.J. Zheng, R.X. Chang, G.Z. Ma, J.M. Xie, Q. Liu, X.R. Liang, and M.H. Ng, Rotavirus infection of the oropharynx and respiratory tract in young children. Journal of medical virology 34 (1991) 29–37.

[11] S. Nishimura, H. Ushijima, S. Nishimura, H. Shiraishi, C. Kanazawa, T. Abe, K. Kaneko, and Y. Fukuyama, Detection of rotavirus in cerebrospinal fluid and blood of patients with convulsions and gastroenteritis by means of the reverse transcription polymerase chain reaction. Brain & development 15 (1993) 457–9.

[12] M. Lynch, B. Lee, P. Azimi, J. Gentsch, C. Glaser, S. Gilliam, H.G. Chang, R. Ward, and R.I. Glass, Rotavirus and central nervous system symptoms: cause or contaminant? Case reports and review. Clinical infectious diseases : an official publication of the Infectious Diseases Society of America 33 (2001) 932–8.

[13] N.G. Feng, A. Sen, M. Wolf, P. Vo, Y. Hoshino, and H.B. Greenberg, Roles of VP4 and NSP1 in Determining the Distinctive Replication Capacities of Simian Rotavirus RRV and Bovine Rotavirus UK in the Mouse Biliary Tract. Journal of virology 85 (2011) 2686–2694.

[14] M. Fenaux, A.A. Cuadras, N. Feng, M. Jaimes, and H.B. Greenberg, Extraintestinal spread and replication of a homologous EC rotavirus strain and a heterologous rhesus rotavirus in BALB/c mice. Journal of virology 80 (2006) 5219–5232.

[15] E.A. Karlsson, V.A. Meliopoulos, C. Savage, B. Livingston, A. Mehle, and S. Schultz-Cherry, Visualizing real-time influenza virus infection, transmission and protection in ferrets. Nature communications 6 (2015) 6378.

[16] N.S. Heaton, V.H. Leyva-Grado, G.S. Tan, D. Eggink, R. Hai, and P. Palese, In vivo bioluminescent imaging of influenza a virus infection and characterization of novel cross-protective monoclonal antibodies. Journal of virology 87 (2013) 8272–81.

[17] V. Tran, L.A. Moser, D.S. Poole, and A. Mehle, Highly sensitive real-time in vivo imaging of an influenza reporter virus reveals dynamics of replication and spread. Journal of virology 87 (2013) 13321–9.

[18] J.F. Rodriguez, D. Rodriguez, J.R. Rodriguez, E.B. McGowan, and M. Esteban, Expression of the firefly luciferase gene in vaccinia virus: a highly sensitive gene marker to follow virus dissemination in tissues of infected animals. Proceedings of the National Academy of Sciences of the United States of America 85 (1988) 1667–71.

[19] B. Manicassamy, S. Manicassamy, A. Belicha-Villanueva, G. Pisanelli, B. Pulendran, and A. Garcia-Sastre, Analysis of in vivo dynamics of influenza virus infection in mice using a GFP reporter virus. Proceedings of the National Academy of Sciences of the United States of America 107 (2010) 11531–11536.

[20] G.D. Luker, J.P. Bardill, J.L. Prior, C.M. Pica, D. Piwnica-Worms, and D.A. Leib, Noninvasive bioluminescence imaging of herpes simplex virus type 1 infection and therapy in living mice. Journal of virology 76 (2002) 12149–61.

[21] S.H. Cook, and D.E. Griffin, Luciferase imaging of a neurotropic viral infection in intact animals. Journal of virology 77 (2003) 5333–8.

[22] C.W. Burke, J.N. Mason, S.L. Surman, B.G. Jones, E. Dalloneau, J.L. Hurwitz, and C.J. Russell, Illumination of parainfluenza virus infection and transmission in living animals reveals a tissue-specific dichotomy. PLoS pathogens 7 (2011) e1002134.

[23] S.S. Gambhir, J.R. Barrio, M.E. Phelps, M. Iyer, M. Namavari, N. Satyamurthy, L. Wu, L.A. Green, E. Bauer, D.C. MacLaren, K. Nguyen, A.J. Berk, S.R. Cherry, and H.R. Herschman, Imaging adenoviral-directed reporter gene expression in living animals with positron emission tomography. Proceedings of the National Academy of Sciences of the United States of America 96 (1999) 2333–2338.

[24] Y. Kanai, S. Komoto, T. Kawagishi, R. Nouda, N. Nagasawa, M. Onishi, Y. Matsuura, K. Taniguchi, and T. Kobayashi, Entirely plasmid-based reverse genetics system for rotaviruses. Proceedings of the National Academy of Sciences of the United States of America 114 (2017) 2349–2354.

[25] S. Komoto, S. Fukuda, T. Ide, N. Ito, M. Sugiyama, T. Yoshikawa, T. Murata, and K. Taniguchi, Generation of Recombinant Rotaviruses Expressing Fluorescent Proteins by Using an Optimized Reverse Genetics System. Journal of virology 92 (2018).

[26] A.A. Philip, J.L. Perry, H.E. Eaton, M. Shmulevitz, J.M. Hyser, and J.T. Patton, Generation of Recombinant Rotavirus Expressing NSP3-UnaG Fusion Protein by a Simplified Reverse Genetics System. Journal of virology 93 (2019).

[27] L. Sanchez-Tacuba, N. Feng, N.J. Meade, K.H. Mellits, P.H. Jais, L.L. Yasukawa, T.K. Resch, B. Jiang, S. Lopez, S. Ding, and H.B. Greenberg, An Optimized Reverse Genetics System Suitable for Efficient Recovery of Simian, Human, and Murine-Like Rotaviruses. Journal of virology 94 (2020).

[28] H. Montero, C.F. Arias, and S. Lopez, Rotavirus Nonstructural Protein NSP3 is not required for viral protein synthesis. Journal of virology 80 (2006) 9031–8.

[29] A.A. Philip, and J.T. Patton, Expression of Separate Heterologous Proteins from the Rotavirus NSP3 Genome Segment Using a Translational 2A Stop-Restart Element. Journal of virology 94 (2020).

[30] S.A. Gonzalez, N.M. Mattion, R. Bellinzoni, and O.R. Burrone, Structure of Rearranged Genome Segment-11 in 2 Different Rotavirus Strains Generated by a Similar Mechanism. Journal of General Virology 70 (1989) 1329–1336.

[31] C.G. England, E.B. Ehlerding, and W. Cai, NanoLuc: A Small Luciferase Is Brightening Up the Field of Bioluminescence. Bioconjugate chemistry 27 (2016) 1175–1187.

[32] C.A. Mebus, and L.E. Newman, Scanning Electron, Light, and Immunofluorescent Microscopy of Intestine of Gnotobiotic Calf Infected with Reovirus-Like Agent. Am J Vet Res 38 (1977) 553–558.

[33] D.R. Snodgrass, K.W. Angus, and E.W. Gray, Rotavirus Infection in Lambs - Pathogenesis and Pathology. Arch Virol 55 (1977) 263–274.

[34] K.W. Theil, E.H. Bohl, R.F. Cross, E.M. Kohler, and A.G. Agnes, Pathogenesis of Porcine Rotaviral Infection in Experimentally Inoculated Gnotobiotic Pigs. Am J Vet Res 39 (1978) 213–220.

[35] J.P. Mcadaragh, M.E. Bergeland, R.C. Meyer, M.W. Johnshoy, I.J. Stotz, D.A. Benfield, and R. Hammer, Pathogenesis of Rotaviral Enteritis in Gnotobiotic Pigs - a Microscopic Study. Am J Vet Res 41 (1980) 1572–1581.

[36] L.M. Little, and J.A. Shadduck, Pathogenesis of Rotavirus Infection in Mice. Infect Immun 38 (1982) 755–763.

[37] O. Oelz, M. Ritter, R. Jenni, M. Maggiorini, U. Waber, P. Vock, and P. Bartsch, Nifedipine for High-Altitude Pulmonary-Edema. Lancet 2 (1989) 1241–1244.

[38] N. Feng, B. Kim, M. Fenaux, H. Nguyen, P. Vo, M.B. Omary, and H.B. Greenberg, Role of interferon in homologous and heterologous rotavirus infection in the intestines and extraintestinal organs of suckling mice. Journal of virology 82 (2008) 7578–7590.

[39] A. Goff, N. Twenhafel, A. Garrison, E. Mucker, J. Lawler, and J. Paragas, In vivo imaging of cidofovir treatment of cowpox virus infection. Virus Res 128 (2007) 88–98.

[40] K. Yamada, K. Noguchi, K. Kimitsuki, R. Kaimori, N. Saito, T. Komeno, N. Nakajima, Y. Furuta, and A. Nishizono, Reevaluation of the efficacy of favipiravir against rabies virus using in vivo imaging analysis. Antivir Res 172 (2019).

[41] D. Pietrella, A. Rachini, A. Torosantucci, P. Chiani, A.J.P. Brown, F. Bistoni, P. Costantino, P. Mosci, C. d’Enfert, R. Rappuoli, A. Cassone, and A. Vecchiarelli, A beta-glucan-conjugate vaccine and anti-beta-glucan antibodies are effective against murine vaginal candidiasis as assessed by a novel in vivo imaging technique. Vaccine 28 (2010) 1717–1725.

[42] U.J. Buchholz, S. Finke, and K.K. Conzelmann, Generation of bovine respiratory syncytial virus (BRSV) from cDNA: BRSV NS2 is not essential for virus replication in tissue culture, and the human RSV leader region acts as a functional BRSV genome promoter. Journal of virology 73 (1999) 251–9.

[43] P.H. Jais, E. Decroly, E. Jacquet, M. Le Boulch, A. Jais, O. Jean-Jean, H. Eaton, P. Ponien, F. Verdier, B. Canard, S. Goncalves, S. Chiron, M. Le Gall, P. Mayeux, and M. Shmulevitz, C3P3-G1: first generation of a eukaryotic artificial cytoplasmic expression system. Nucleic acids research 47 (2019) 2681–2698.

[44] L. Sanchez-Tacuba, M. Rojas, C.F. Arias, and S. Lopez, Rotavirus Controls Activation of the 2’-5’-Oligoadenylate Synthetase/RNase L Pathway Using at Least Two Distinct Mechanisms. Journal of virology 89 (2015) 12145–53.

